# *Park7* deletion leads to age- and sex-specific transcriptome changes involving NRF2-CYP1B1 axis in mouse midbrain astrocytes

**DOI:** 10.1101/2024.02.23.581743

**Authors:** Sergio Helgueta, Tony Heurtaux, Alessia Sciortino, Yujuan Gui, Jochen Ohnmacht, Pauline Mencke, Ibrahim Boussaad, Rashi Halder, Pierre Garcia, Rejko Krüger, Michel Mittelbronn, Manuel Buttini, Thomas Sauter, Lasse Sinkkonen

**Affiliations:** Department of Life Sciences and Medicine (DLSM), University of Luxembourg, Belvaux, Luxembourg; Luxembourg Centre of Neuropathology (LCNP), Luxembourg; Luxembourg Centre for Systems Biomedicine (LCSB), University of Luxembourg, Belvaux, Luxembourg; Centre Hospitalier de Luxembourg (CHL), L-1210, Luxembourg, Luxembourg; Luxembourg Institute of Health (LIH), L-1445, Luxembourg, Luxembourg; National Center of Pathology (NCP), Laboratoire National de Santé (LNS), Dudelange, Luxembourg

**Keywords:** *Park7*, transcriptomics, sexual dimorphism, astrocytes, oxidative stress, NRF2

## Abstract

Loss-of-function mutations in *PARK7*, encoding for DJ-1, can lead to early onset Parkinson’s disease (PD). In mice, *Park7* deletion leads to dopaminergic deficits during aging, and increased sensitivity to oxidative stress. However, the severity of the reported phenotypes varies. To understand the early molecular changes upon loss of DJ-1, we performed transcriptomic profiling of midbrain sections from young mice. Interestingly, while at 3 months the transcriptomes of both male and female mice were unchanged compared to their wildtype littermates, an extensive deregulation was observed specifically in 8-month-old males. The affected genes are involved in processes such as focal adhesion, extracellular matrix interaction, and epithelial-to-mesenchymal transition (EMT), and enriched for primary target genes of Nuclear factor erythroid 2-related factor 2 (NRF2). Consistently, the antioxidant response was altered specifically in the midbrain of male DJ-1 deficient mice. Many of the misregulated genes are known target genes of estrogen and retinoic acid signaling and show sex-specific expression in wildtype mice. Depletion of DJ-1 or NRF2 in male, but not female primary astrocytes recapitulated many of the *in vivo* changes, including downregulation of cytochrome P450 family 1 subfamily B member 1 (CYP1B1), an enzyme involved in estrogen and retinoic acid metabolism. Interestingly, knock-down of CYP1B1 led to gene expression changes in focal adhesion and EMT in primary male astrocytes. Finally, male iPSC-derived astrocytes with loss of function mutation in the *PARK7* gene also showed changes in the EMT pathway and NRF2 target genes. Taken together, our data indicate that loss of *Park7* leads to sex-specific gene expression changes specifically in males through astrocytic alterations in the NRF2-CYP1B1 axis. These findings suggest higher sensitivity of males to loss of DJ-1 and might help to better understand variation in the reported *Park7^−/−^*phenotypes.

## Introduction

The clinical diagnosis of Parkinson’s disease (PD) relies on the presence of specific motor symptoms, although non-motor symptoms such as hyposmia, depression, anxiety, dementia, autonomic and cognitive dysfunction or REM sleep behavior disorder can be present up to 20 years prior to initial PD diagnosis ^1,2^. PD has an increasing incidence in the population above the age of 65 ^3^, while prevalence and incidence of PD are higher in males than in females ^4,5^. One of the most characteristic neuropathological hallmarks of PD is the progressive loss of dopaminergic neurons in the substantia nigra pars compacta (SNpc of the midbrain) leading to striatal dopamine deficiency ^6^. One of the main drivers of this loss is oxidative stress and subsequent mitochondrial dysfunction, resulting in increased levels of reactive oxygen species (ROS) ^7–9^. Neurons depend on glial cells such as microglia, oligodendrocytes, and astrocytes for their protection against oxidative stress. The accumulation of ROS in all these cell types is sensed by NRF2 pathway through oxidation of Kelch-like ECH-associated protein 1 (KEAP1) that leads to release and stabilization of Nuclear factor erythroid 2-related factor 2 (NFE2L2, also known as NRF2), and its nuclear translocation, thereby inducing the expression of antioxidant response genes. A failure in antioxidant response can lead to oxidative stress ^10^. Interestingly, the antioxidant response can be modulated by developmental signaling pathways and sexual hormones such as retinoic acid and estrogen, respectively, leading to differences in response depending on the cell state or sex ^11^. In particular estrogen is associated with neuroprotection, possibly due to improved antioxidant response, and could contribute to sexual dimorphism of disease-risk in many disorders, including PD ^12,13^

DJ-1 (encoded by *Parkinsonism associated deglycase* or *PARK7*) has been characterized as a causative gene for autosomal recessive, early-onset PD ^14^ and the PD-associated mutations in the gene are typically leading to loss or reduction of the functional DJ-1 protein ^15^. To date, various functions have been reported for DJ-1, mostly associated with mitochondrial biology and oxidative stress ^16^. In both mouse and human cell lines, DJ-1 was shown to stabilize NRF2 through inhibition of its interaction with KEAP1^17^. Different primary mouse cell and animal models have been utilized to investigate the consequences of *Park7* gene disruption. However, the literature shows inconsistent results with respect to *Park7^−/−^* mouse models for studying PD. Most of the studies show no dopaminergic neurodegeneration in the SNpc ^18–22^. In addition, Pham and colleagues show that these mice have less dopaminergic neurons in the VTA and exhibit non-motor symptoms associated with early phases of PD, such as impairment in motivated behavior and cognition ^22^. Goldberg et al. reported abnormal nigral dopaminergic physiology in *Park7^−/−^* mice, including altered excitability and firing of dopamine neurons projecting to the striatum, together with defects in locomotor activity ^18^. In contrast to these studies, Rousseaux and colleagues report that their *Park7*^−/−^ mouse model show early-onset progressive dopaminergic cell loss, advancing from unilateral to bilateral with aging ^23^. The authors also see degeneration at the locus coeruleus and mild motor behavior deficits at later time points ^23^. Taken together, controversy still exists regarding the dopaminergic system phenotype exhibited by *Park7^−/−^* mice, with notable variations across studies. It is worth noting that some previous studies did not specify the sex of the used animals, which is a critical variable that could influence the observed phenotypes in this mouse model.

Here, we investigated the early gene expression changes in the midbrain of both male and female *Park7^−/−^* mice to obtain insights into the early events triggered by absence of DJ-1 *in vivo* and to investigate the possible sex-specific differences. Moreover, we performed cell type-specific investigation of *Park7*-depletion in cultured primary astrocytes through RNAi experiments, and confirmed the disease-relevance of our findings using human iPSC-derived astrocytes carrying a *PARK7* mutation. Together our results identify many age-dependent changes specifically in the male midbrain with processes such as EMT and focal adhesion becoming altered in astrocytes in an NRF2-dependent manner. These findings could have implications on understanding of the mechanisms of DJ-1-associated PD and the higher PD-risk of males, and help to reconcile discrepant results regarding different *Park7^−/−^*-mouse models.

## Results

### *Park7* deletion induces transcriptomic changes specifically in midbrain of 8-month-old males

To identify the early gene expression changes induced by loss of DJ-1 in ventral midbrain in both males and females, we dissected midbrains from 3- and 8-month-old *Park7^−/−^*and wildtype littermate control mice and isolated total RNA for RNA sequencing (RNA-seq) (Fig. 1). Interestingly, only 3 and 7 differentially expressed genes (DEGs, FDR < 0.05) were identified in females and males, respectively, at the age of 3 months (Fig. 1b-c, Supplementary Table 1-2). In contrast, by 8 months the number of DEGs in females increased to 101 (Fig. 1d) and in males over 1200 DEGs with generally stronger fold changes were detected (Supplementary Table 1-2). To confirm the extensive changes observed in males, we collected additional midbrain samples for RNA-seq analysis from an independent cohort of mice, with many of the gene expression changes remaining (Supplementary Table 1-2). After combining the independent cohorts and applying a batch correction, 746 mostly downregulated DEGs were confirmed in males (Fig. 1e, Supplementary Table 1). Strikingly, only 8 of the 746 DEGs in males were shared with the 101 DEGs identified in females at the same age. Finally, to confirm that the lack of changes in 8-month-old females is not due to technical or biological variation between the female samples, for example due to oestrus cycle, we analyzed the per-gene dispersions to estimate the median dispersion of each sample set (Supplementary Figure 1). Interestingly, the second cohort of 8-month-old male samples showed increased dispersion compared to other sample sets, partially explaining the lower number of DEGs compared to the first cohort. However, importantly, the median dispersion in 8 month old females was not higher compared males. Thus, our results indicate that loss of DJ-1 leads to age- and sex-specific gene expression changes in the midbrain, with males particularly affected.

**Figure 1:**
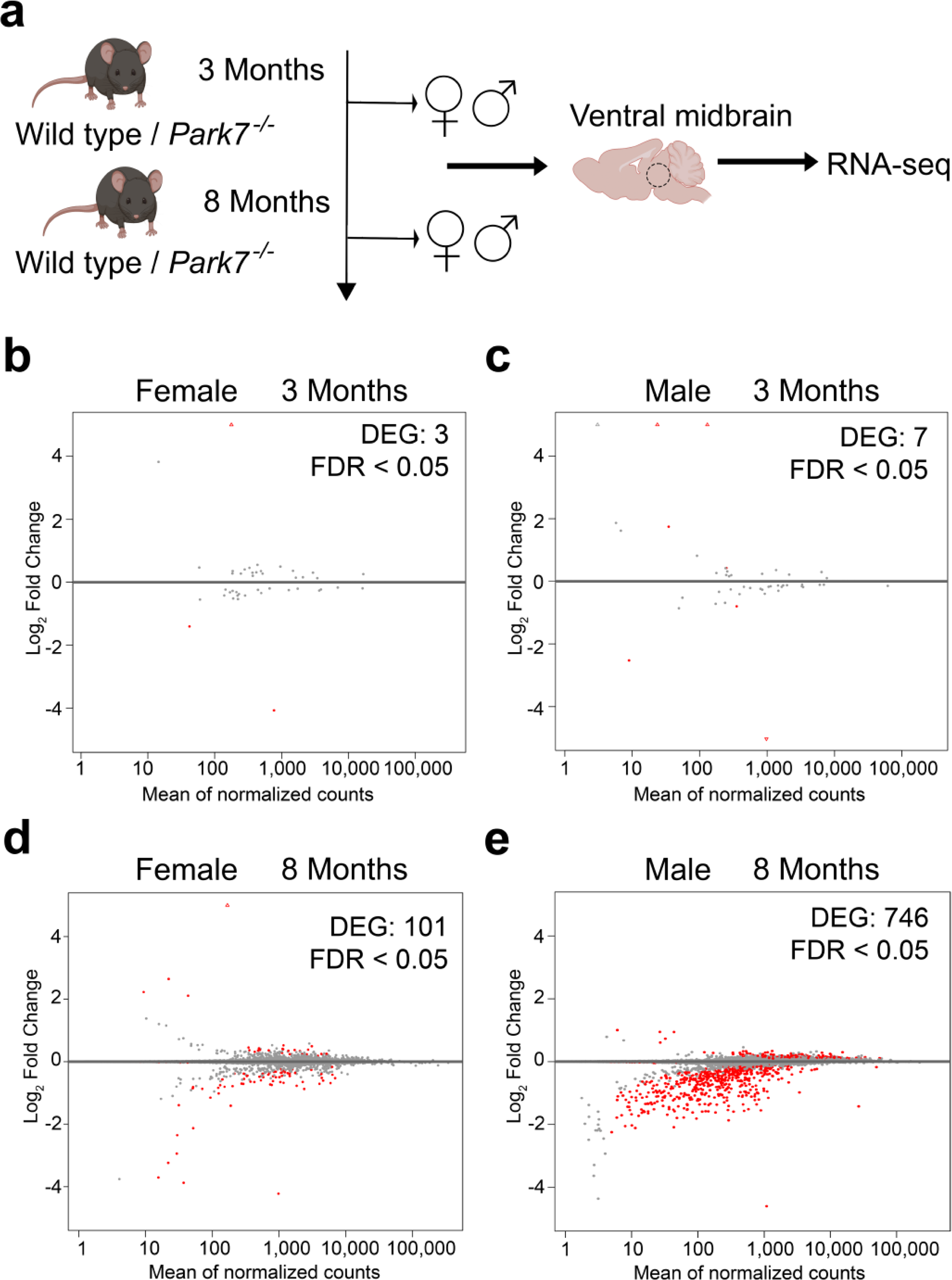
*Park7* deletion induces transcriptome deregulation during ageing predominantly in male ventral midbrain. **(a)** Schematic diagram of the experimental set-up for genome-wide transcriptomics profiling of isolated midbrains of 3-month-old and 8-month-old wild type and *Park7^−/−^*mice. **(b-d)** MA plots showing DEGs in the midbrains of *Park7*^−/−^ mice compared to wildtype littermate controls in four different age- and sex-groups, namely **(b)** 3-month-old females (3 DEGs, n=4), (**c)** 3-month-old males (7 DEGs, n=4), **(d)** 8-month-old females (101 DEGs, n=4), and **(e)** 8-month-old males (746 DEGs, n=8). Each red dot represents one DEG defined by FDR < 0.05. The x-axis represents the mean of normalized counts and the y-axis the log_2_-fold change. Full lists of DEGs are available in Supplementary Table S1.

### Loss of DJ-1 in male mice downregulates genes involved in epithelial to mesenchymal transition, focal adhesion, and extracellular matrix composition

To better understand the male-specific gene regulatory changes, we performed enrichment analysis for the identified 746 DEGs. Focal adhesion was found to be the most enriched pathway in KEGG and WikiPathways databases and also among the top 5 GO cellular components (Fig. 2a-d, Supplementary Table 3). Consistently, extracellular matrix (ECM)-receptor interaction and regulation of ECM were found among the top pathways, indicating changes in cellular adhesion and integrin-mediated interaction with ECM in the *Park7^−/−^* mice. Other top pathways included PI3K-Akt signaling pathway, interferon gamma response, and early estrogen response. Finally, the process with most significant enrichment across the different databases was found to be epithelial mesenchymal transition (EMT) in the MSigDB Hallmark database (Fig. 2c). Indeed, 58 genes associated with EMT were significantly differentially expressed, with almost all of the genes showing downregulation in *Park7^−/−^* mice (Fig. 2e). Consistenly, *Cdh1*, encoding for E-cadherin, a well established marker of EMT ^24^, was the gene with strongest fold change across all of the DEGs (Fig. 2f, Supplementary Table 1). *Twist1*, encoding for a transcriptional regulator typically induced in EMT, was also among the strongly downregulated genes, indicating that the cells are not undergoing classical EMT but rather an EMT-like process involving many of the same genes. Importantly, the downregulation of *Cdh1* was confirmed by real time quantitative PCR (RT-qPCR) to take place in midbrain of 8-month-old males, but was not observed in females or in isolated cortex from the same male mice (Fig. 2g). Enrichment analysis for the 101 DEGs obtained for 8-month-old females did not show any of the enriched pathways from males (Supplementary Figure 2a-d, Supplementary Table 4)

**Figure 2:**
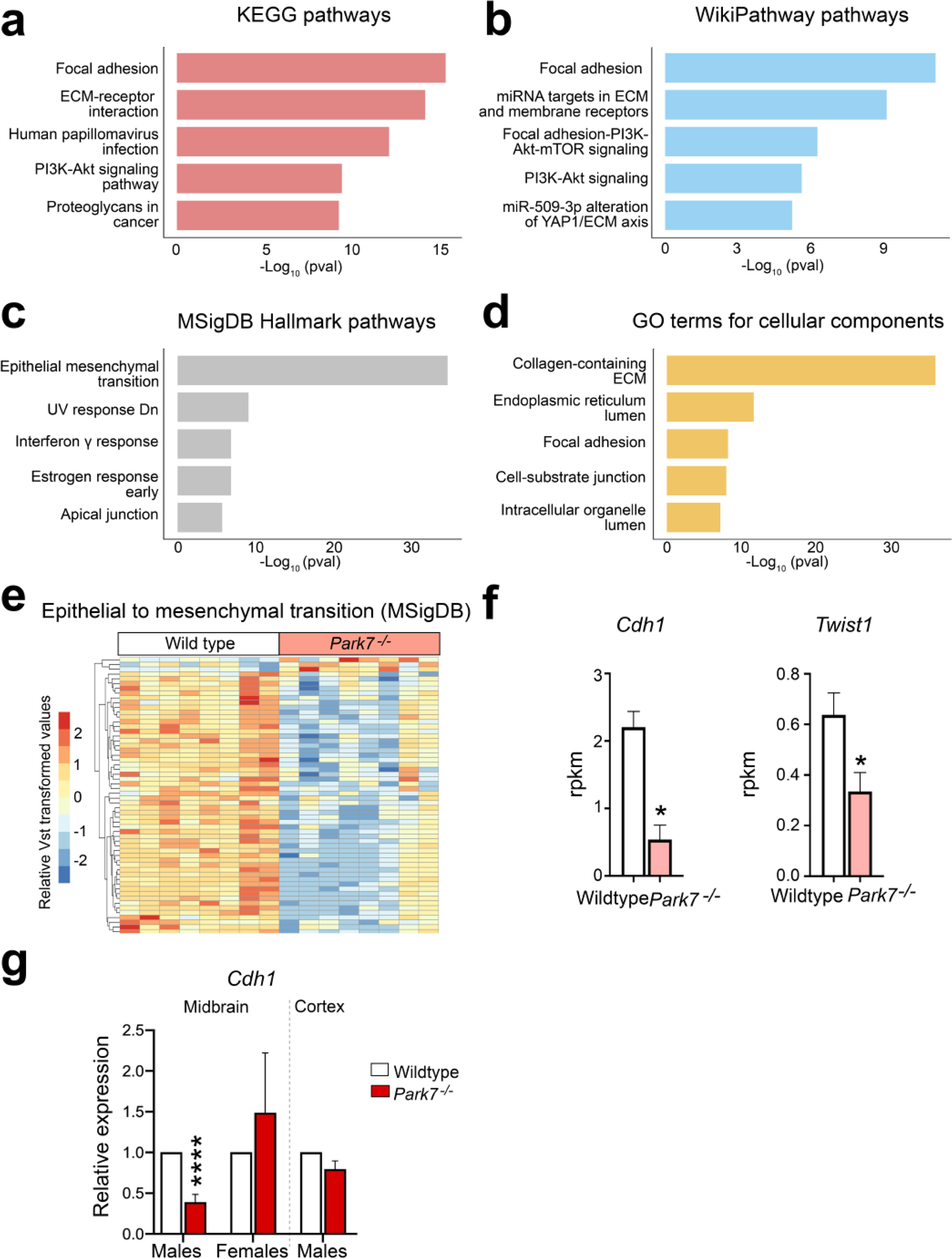
Loss of *Park7* alters expression of genes involved in focal adhesion, epithelial-to-mesenchymal transition, and extracellular matrix composition only in midbrain. **(a-d)** Pathway enrichment analysis for the 746 DEGs altered in 8-month-old male mice. The top 5 results from **(a)** KEGG pathways, (**b)** WikiPathway pathways, **(c)** MSigDB Hallmark pathways, and **(d)** GO terms for cellular components based on the significance of enrichment are shown.The x-axis represents the -log_10_ p-value of pathway enrichment. Complete results of enrichment analysis are available in the Supplementary Table S2. **(e)** Heatmap showing the relative expression of the 58 genes associated with EMT and found to be differentially expressed in midbrains of male *Park7*^−/−^ mice compared to wild type littermates at 8-months of age. Vst-transformed read counts are plotted with each row representing one gene and each column representing an independent midbrain sample. **(f)** Expression levels in RPKM obtained by RNA-seq showing significant downregulation in the expression of *Cdh1* and *Twist1* in the midbrain of *Park7*^−/−^ male mice at 8-months of age in comparison to wild type mice. * = FDR < 0.05. (**g)** RT-qPCR validation of *Cdh1* downregulation in the midbrain of *Park7*^−/−^ male mice at 8-months of age. No downregulation was observed in female midbrain or in male cortex. Values represent mean ± SEM (n=4-8 mice per group). Statistical significance was tested by unpaired t-test. **** = p-value < 0.0001.

### Sex-specific changes in *Park7^−/−^* mice are associated with lowered NRF2 signaling

Next we asked how the loss of DJ-1 can lead to the observed transcriptional changes and what are the transcription factors (TFs) involved. To address this, we took advantage of the existing public data on TF target genes identified through chromatin immunoprecipitation (ChIP) experiments and derived from the ENCODE project and ChIP Enrichment Analysis (ChEA) database ^25^. When focusing on bona fide TFs, the strongest enrichment among the male, but not female, DEGs was found for the primary targets of NRF2 (encoded by *Nfe2l2* gene) (Fig. 3a) (Supplementary Figure 2e) These consisted of over 70 DEGs that are known primary NRF2 targets, most of which were downregulated and found to be associated with EMT process (Fig. 3b). However, some known NRF2 target genes involved in reactive oxygen species (ROS) production were also found to be upregulated. These results indicate that NRF2 could be mediating the DJ-1 induced transcriptional changes, involving an overall decrease in NRF2-mediated signalling and antioxidant response. Indeed, analysis of well established antioxidant response genes confirmed their downregulation in *Park7^−/−^* mice (Fig. 3c). *Nfe2l2* itself also showed a trend of downregulation but was not found to be significant after multiple testing correction (FDR = 0.144). Importantly, for representative antioxidant response genes *Gpx8*, *Gstm2*, and *Ephx1,* the downregulation in male midbrain was again confirmed by RT-qPCR and found to be absent or modest in females or in cortex of the same males (Fig. 3d-g). Interestingly, by RT-qPCR also the downregulation of *Nfe2l2* was found to be significant in male midbrain (Fig. 3d), consistent with the known autoregulation of *Nfe2l2* locus by NRF2 ^26^.

**Figure 3:**
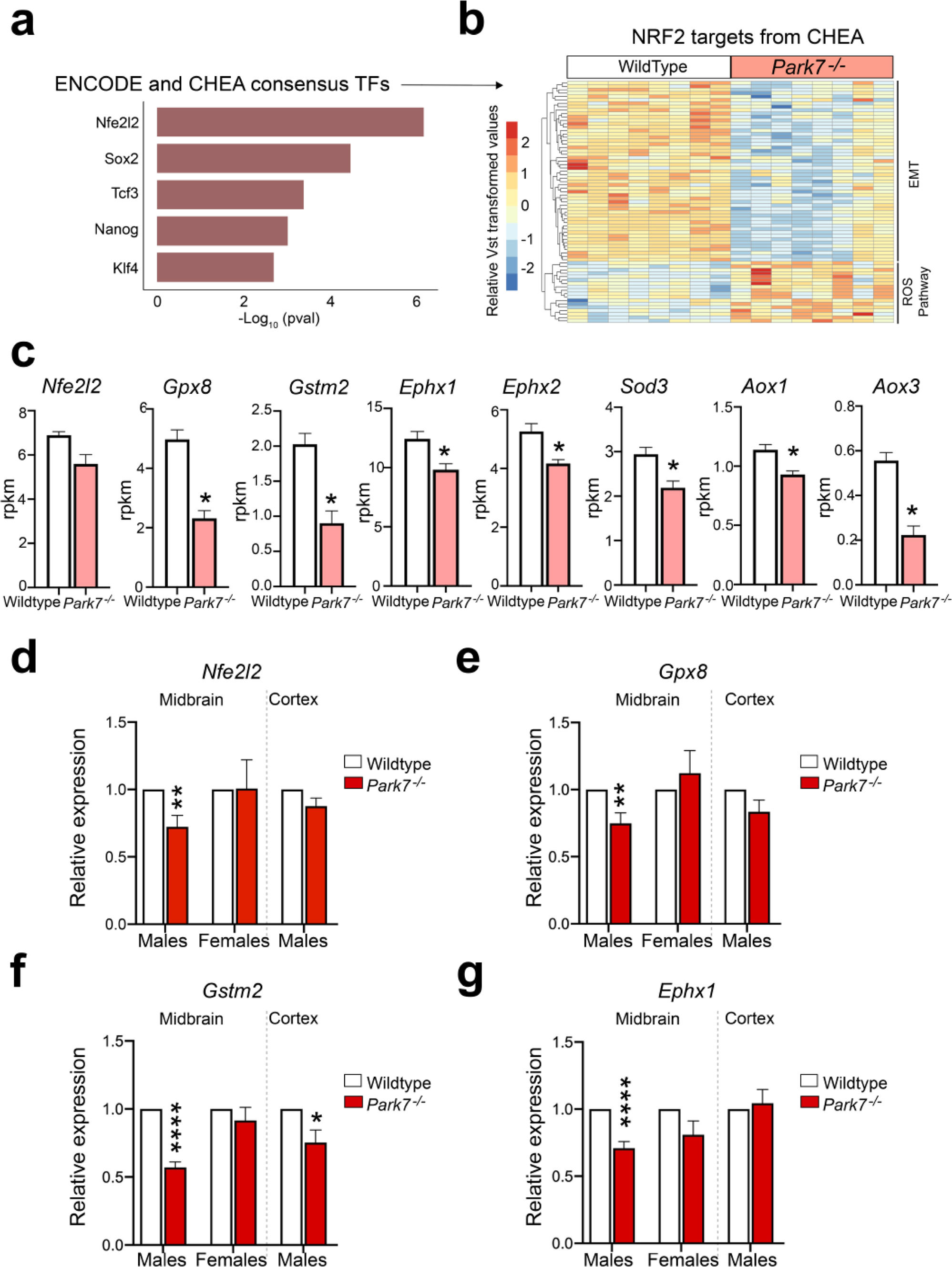
*Park7* depletion leads to reduced antioxidant response in mouse midbrain. **(a)** Top 5 TFs associated with DEGs from *Park7*^−/−^ male mice based on the significance of enrichment for primary TF targets from the ENCODE project and ChEA database. NRF2, encoded by *Nfe2l2* gene, is the most enriched TF. The x-axis represents the -log_10_ p-value. **(b)** Heatmap showing the relative expression of the 70 primary NRF2 target genes from CHEA database found to be differentially expressed in midbrains of male *Park7*^−/−^ mice compared to wild type littermates at 8-months of age. Vst-transformed read counts are plotted with each row representing one gene and each column representing an independent midbrain sample. **(c)** Expression levels in RPKM obtained by RNA-seq showing significant downregulation in the expression of several antioxidant response genes in the midbrain of *Park7*^−/−^ male mice at 8-months of age in comparison to wild type mice. * = FDR < 0.05. (**d-g)** RT-qPCR validation of **(d)** *Nfe2l2,* **(e)** *Gpx8,* **(f)** *Gstm2*, and **(g)** *Ephx1* downregulation in the midbrain of *Park7*^−/−^ male mice at 8-monts of age. No downregulation was observed in female midbrain or in male cortex except for *Gstm2*. Values represent mean ± SEM (n=4-8 mice per group). Statistical significance was tested by unpaired t-test. * = p-value < 0.05; ** = p-value < 0.01; **** = p-value < 0.0001.

### Genes altered by *Park7* deletion show sex-specific expression

To extend the analysis of the upstream events beyond direct TF binding, we performed upstream regulator prediction using Ingenuity Pathway Analysis (IPA) tool. Interestingly, the top upstream regulator predicted to explain most of expression changes in males was beta-estradiol, also known as *17β*-oestradiol, a key sex hormone and agonist for TFs called estrogen receptors (ERs) (Fig. 4a) ^27^. This was not observed for females (Supplementary Figure 2f). The top 5 regulators of male DEGs also included TGFβ, Agt, lipopolysaccharide, and tretinoin, also known as *all-trans* retinoic acid, an agonist of retinoic acid receptors (RARs). ERs and RARs have been described to exert opposing effects on their shared target genes through genomic antagonism ^28^, while both pathways are also known to influence NRF2 signaling ^10^.

**Figure 4.**
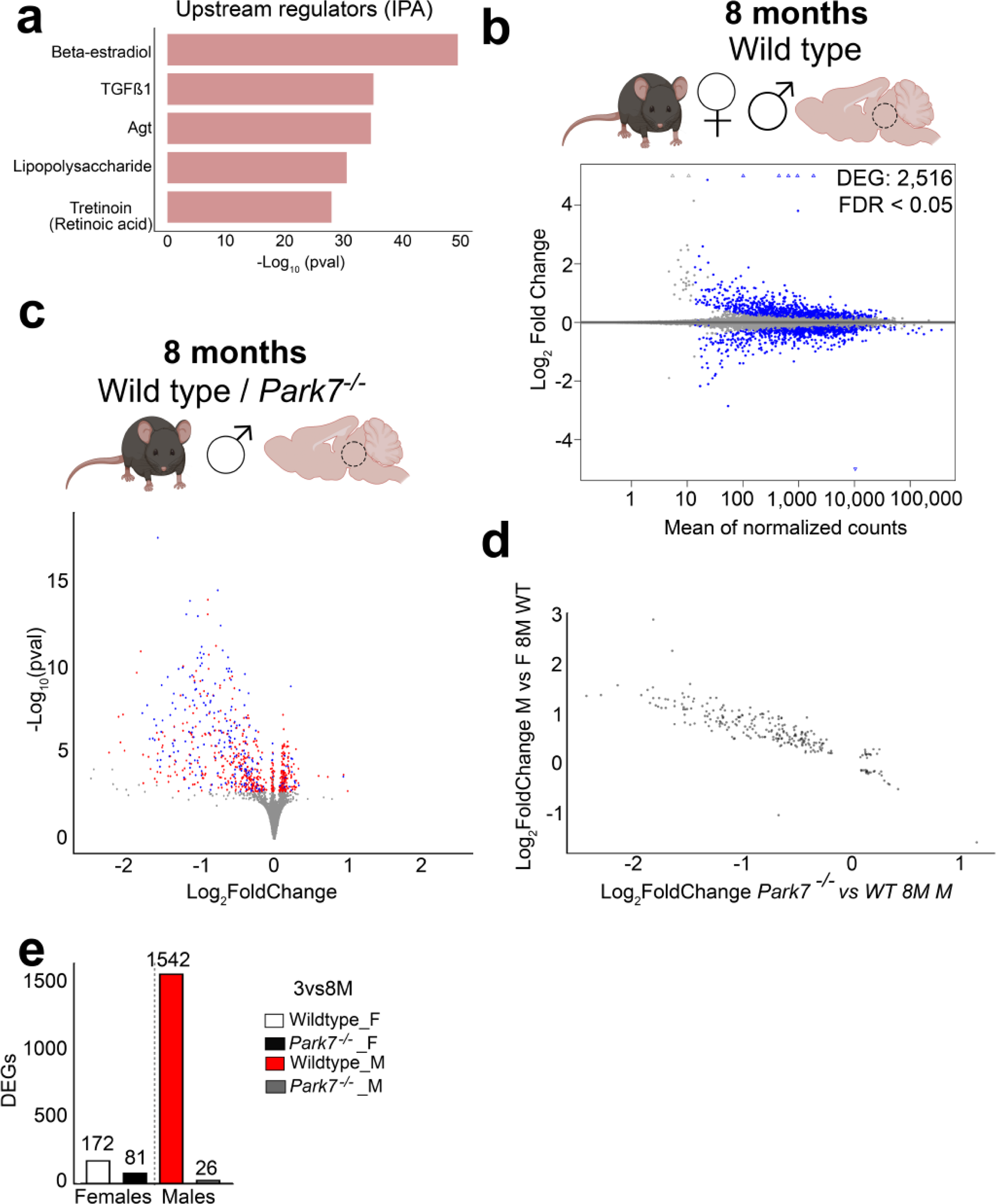
*Park7* deletion affects genes with sex-specific expression profile in midbrain. **(a)** Top5 Upstream Regulators associated with DEGs from *Park7*^−/−^ male mice predicted by Ingenuity Pathway Analysis (IPA). The x-axis represents the -log_10_ p-value of pathway enrichment. Beta-estradiol is the most enriched regulator while retinoic acid (tretinoin) is fifth most significant. **(b)** MA plot depicting the 2516 DEGs between wild type male and female midbrain in 8-month-old mice (n=4-8). Each blue dot represents a DEG defined by FDR < 0.05. The x-axis represents the mean of normalized counts and the y-axis the log_2_-fold change. **(c)** Volcano plot depicting the 746 DEGs altered in 8-month-old male mice and their overlap with the 2516 DEGs from the panel b (n=8). Each red dot represents a DEG only upon *Park7*-depletion (FDR < 0.05) while each blue dot represents a DEG in both *Park7*-depletion and in comparison between wild type males and females (FDR < 0.05). Hypergeometric p-value of the overlap is 0.0069. The x-axis represents the log_2_-fold change and y-axis the -log_10_ p-value. **(d)** Correlation plot of the log_2_-fold changes of the overlapping genes in the two different comparisons, revealing a negative correlation. The x-axis represents the log_2_-fold change of the pairwise comparison between *Park7^−/−^* and wild type 8-month-old male mice. The y-axis represents the log2-fold change of the same gene for pairwise comparison between wild type male and female mice. **(e)** The number of genes with differential expression during aging from 3 months to 8 months per genotype and sex.

Since the midbrain gene expression changes upon DJ-1 depletion are exhibiting sexual dimorphism, and oestradiol was recently described to control sex differences in the brain ^29^, we set out to identify the sexually dimorphic genes in the mouse midbrain at baseline. Differential gene expression analysis of the 8-month-old wildtype male and female midbrains identified 2516 DEGs between the sexes (Fig. 4b). Comparison of these genes with those misregulated in *Park7^−/−^* male mice showed an extensive and significant overlap, indicating that a large proportion of genes affected by loss of DJ-1 are also differentially expressed between males and females at baseline (Fig. 4c). Indeed, comparison of the fold changes of the overlapping genes in the different comparisons revealed a negative correlation, suggesting that genes with normally higher expression in males are reduced in expression in *Park7^−/−^*male mice (Fig. 4d).

To investigate this observation as a function of aging, we compared the DEG’s expression changes between 3 and 8 months for each strain and sex. Strikingly, while the affected genes showed very few changes during aging in female mice or in *Park7^−/−^* males, a strong induction in expression could be observed for wildtype male mice (Figure 4e). Suggesting that in wildtype males a sex-specifc expression increase occurs during aging for genes involved in processes such as anti-oxidant response. And this increase is not observed in absence of DJ-1.

### DJ-1-NRF2-CYP1B1 axis regulates *Cdh1* expression in mouse astrocytes

Among the genes with higher expression in males and significant reduction in *Park7^−/−^* mice, we identified *Cyp1b1*, encoding a cytochrome P450 family enzyme that is involved in the metabolism of both *17β*-oestradiol and *all-trans* retinoic acid (Fig. 5a) ^30,31^. *Cyp1b1* was downregulated around 2-fold in the male midbrain as measured by both RNA-seq and RT-qPCR and, similarly to other NRF2 target genes ^32^, this downregulation was not observed in females or in male cortex (Fig. 5b). The expression level of *Cyp1b1* in the midbrain samples was relatively low (approximately 1 read per kilo per million (RPKM)), suggesting that *Cyp1b1* might be selectively expressed in only some of the cell types of the brain. Indeed, analysis of Brain RNA-seq database ^33^ for purified cell types from mouse brain confirmed *Cyp1b1* to be mostly expressed in mouse astrocytes (Fig. 5c).

**Figure 5.**
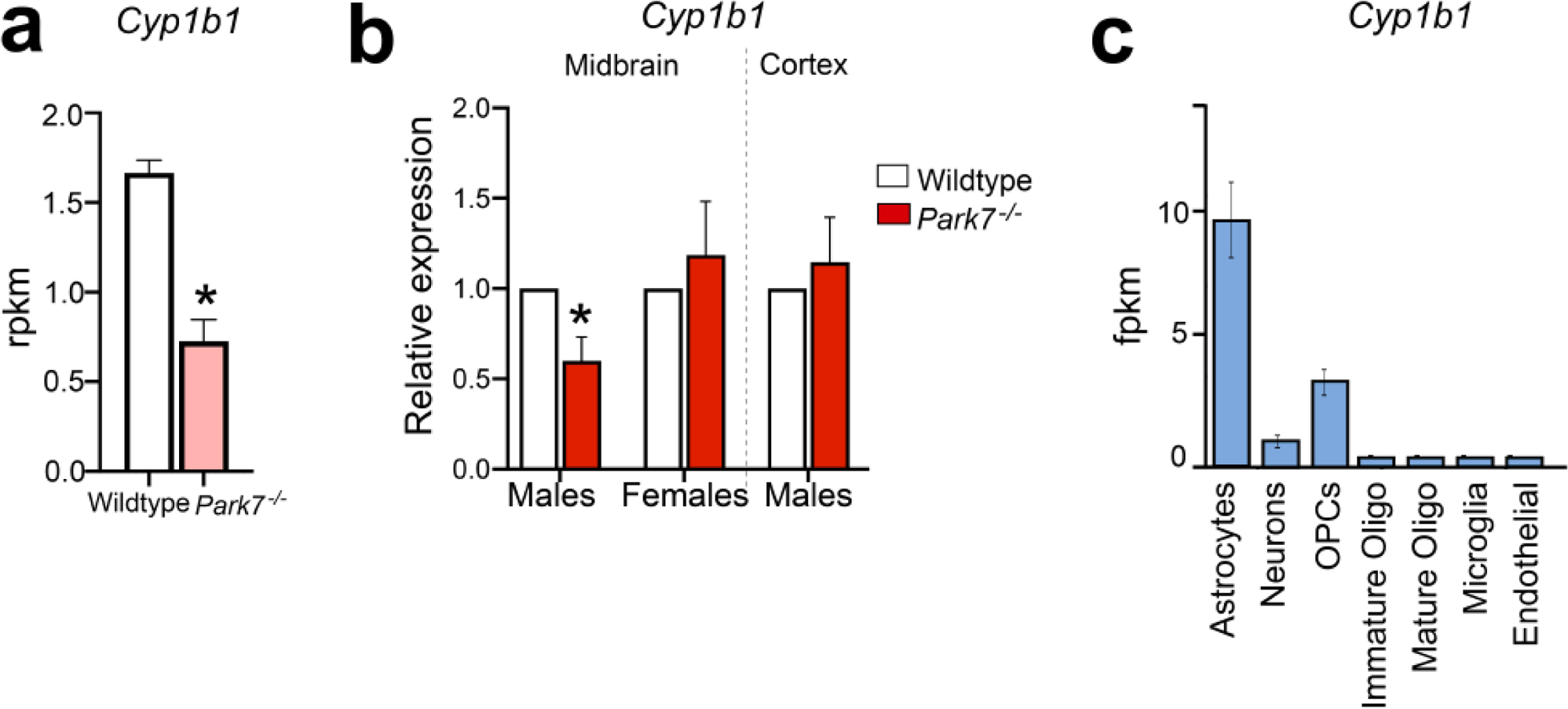
Downregulation of Cyp1b1 in the midbrain of male *Park7^−/−^* mice. **(a)** Expression levels in RPKM obtained by RNA-seq showing significant downregulation in the expression of *Cyp1b1* in the midbrain of *Park7*^−/−^ male mice at 8-months of age in comparison to wild type mice. * = FDR < 0.05. **(b)** RT-qPCR validation of *Cyp1b1* downregulation in the midbrain of *Park7*^−/−^ male mice at 8-months of age. No downregulation was observed in female ventral midbrain or in male cortex. Values represent mean ± SEM (n=4-8 mice per group). Statistical significance was tested by unpaired t-test. * = p-value < 0.05. **(c)** Expression levels as fragments per kilobase million (FPKM) of *Cyp1b1* in the different cell populations of the mouse brain obtained from the Brain RNA-seq database from Barres’ lab (http://www.brainrnaseq.org/). *Cyp1b1* is mostly expressed in mouse astrocyte.

To analyse the possible contribution of *Cyp1b1* downregulation to the observed expression changes of genes involved in processes such as EMT, we isolated primary astrocytes separately from newborn male and female mice for RNA interference (RNAi) experiments (Fig. 6a). In parallel, microglia and oligodendrocyte precursors were also isolated as additional cell types with highly active NRF2 pathway but with lower *Cyp1b1* expression. The identities of the isolated cells were confirmed by stainings for markers of astrocytes (GFAP), microglia (F4/80), and oligodendrocytes (MBP) (Fig. 6b), and the sex of the mice was confirmed by PCR genotyping using *Ube* primers (Supplementary Figure 3). Following the transfection of siRNAs targeting *Park7* mRNA, a strong downregulation of *Park7* was observed in all three cell types (Fig. 6c). Interestingly, this led to a significant downregulation of *Nfe2l2*, *Cyp1b1*, and the EMT-marker *Cdh1* in male astrocytes, but not in microglia and only for *Cyp1b1* in oligodendrocytes. Moreover, a downregulation of *Cdh1* was also observed in female astrocytes while an unexpected upregulation was detected in female microglia. To test whether the effect on *Cdh1* was mediated via NRF2 signaling, we also performed an *Nfe2l2* knock-down (Fig. 6d). Consistently, *Cyp1b1* and *Cdh1* were downregulated in male astrocytes, and for *Cyp1b1* also in female astrocytes, while microglia and oligodendrocytes remained largely unaffected. Finally, to test whether the misregulation of *Cyp1b1* could contribute to changes in *Cdh1* expression, we transfected oligodendrocytes, microglia and astrocytes with siRNAs targeting *Cyp1b1* and evaluated the expression of genes along the DJ-1-NRF2-CYP1B1 axis. While *Park7* and *Nfe2l2* were not reduced by *Cyp1b1* depletion, confirming their function upstream of *Cyp1b1*, *Cdh1* was significantly downregulated, especially in male astrocytes (Fig. 6e). Thus, the EMT pathway alterations in the midbrain of *Park7^−/−^* male mice are likely to take place in astrocytes and involve the NRF2-CYP1B1 axis.

**Figure 6.**
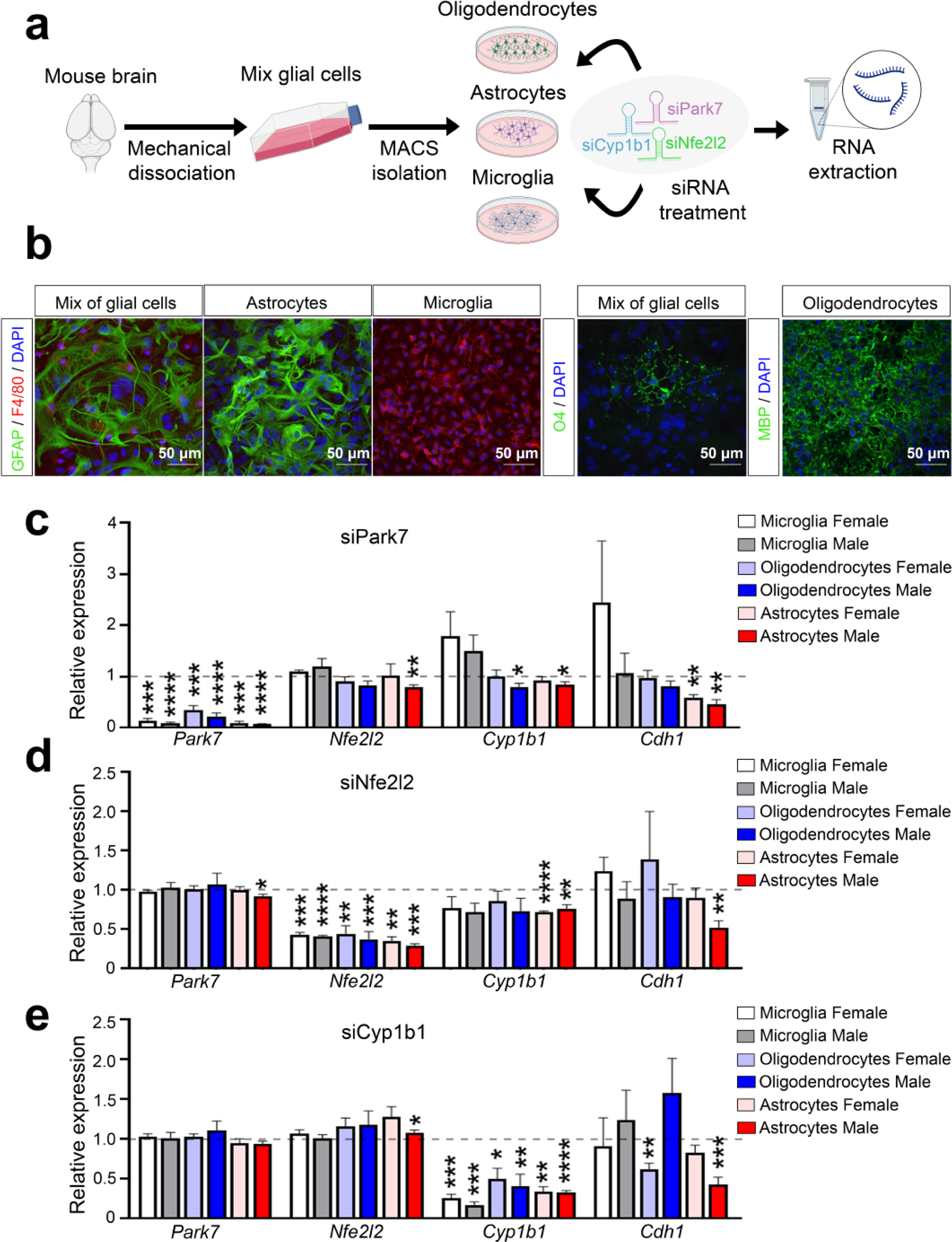
Depletion of *Park7*, *Nfe2l2*, and *Cyp1b1* induces downregulation of *Cdh1* in primary male astrocytes but not in microglia or oligodendrocytes. **(a)** Schematic diagram of the experimental set-up for glial cell isolation from newborn mice followed by RNAi experiments. Mix of glial cells of each sex were cultured separately after brain removal and mechanical dissociation. From the mixture of glial cells, microglia, astrocytes, and oligodendrocytes were isolated using magnetic cell sorting (MACS) and cultured up to 2 weeks prior to siRNA transfection and RNA extraction 24 h post-transfection. **(b)** Identity of the isolated cells was confirmed by stainings for GFAP (astrocyte marker, green), F4/80 (microglia marker, red), O4 and MBP (oligodendrocyte markers, green), and DAPI in blue as the nuclear marker. **(c-e)** RT-qPCR of gene expression changes upon transfection of siRNAs targeting **(c)** *Park7*, **(d)** *Nfe2l2*, or **(e)** *Cyp1b1*. Expression of *Park7*, *Nfe2l2*, *Cyp1b1* and *Cdh1* was normalized to siScramble per replicate (indicated by dashed line). Values represent mean ± SEM (n = 3-7 mice per group). Statistics were tested by one sample t-test, taking 1 as the theoretical mean. * = p-value < 0.05, ** = p-value < 0.01, *** = p-value < 0.001, and **** = p-value < 0.0001.

### *Park7* deletion induced changes in focal adhesion and epithelial to mesenchymal transition involve NRF2 and CYP1B1 in astrocytes

To evaluate the genome-wide impact of the different knock-downs, we performed transcriptome analysis of the male astrocytes 24 hours post-transfection and compared to cells similarly treated with an unspecific control siRNA. Knock-down of *Park7*, *Nfe2l2*, and *Cyp1b1* induced a significant differential expression (FDR < 0.05) of 673, 1055, and 918 genes, respectively, when performing paired analysis (Fig. 7a-c, Supplementary Table 5). Also in primary astrocytes, more down-than upregulated genes were observed for each knock-down. Many of the DEGs were different from those detected in isolated midbrain, revealing the limitations of cell culture systems in mimicking the adult brain in the absence of the appropriate microenvironment. Interestingly, the upregulated genes were enriched for pathways such as interferon signaling and primary targets of IRF8 (Supplementary Figure 4). However, when focusing on the top 5 pathway enrichments for downregulated genes upon *Park7* depletion in MSigDB Hallmark and WikiPathways databases, EMT and focal adhesion were among the top enriched pathways also in cultured astrocytes (Fig. 7d-e). When analysing the enrichments of the same top 5 pathways among the genes downregulated upon *Nfe2l2* or *Cyp1b1* depletion, it was clear that many of the pathways altered by *Park7* depletion, such as PI3K-Akt and interferon signaling, were not similarly affected (Fig. 7e and Supplementary Fig. 4). But importantly, both EMT and focal adhesion were significantly enriched also in these conditions. Suggesting that *Park7* deletion induced gene expression changes affecting EMT and focal adhesion are occurring in astrocytes and are mediated by NRF2 and CYP1B1. In keeping with this result, the downregulated genes for each knock-down in astrocytes were enriched for primary target genes of NRF2 (Fig. 7f).

**Figure 7.**
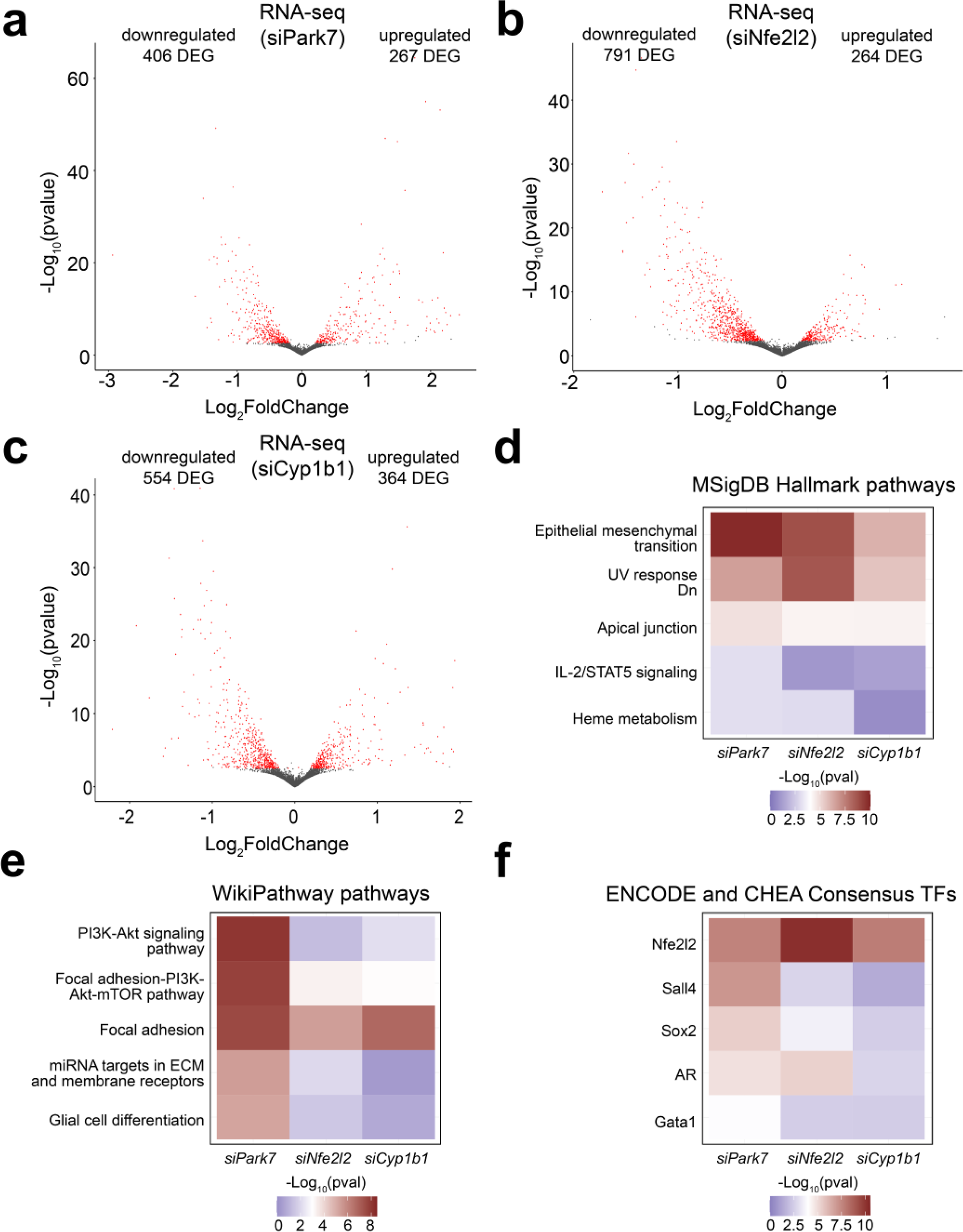
Depletion of each *Park7*, *Nfe2l2*, and *Cyp1b1* in primary astrocytes induces downregulation of genes involved in focal adhesion, epithelial-to-mesenchymal transition, and extracellular matrix composition. **(a-c)** Volcano plot depicting the DEGs in primary mouse astrocytes upon the knockdown of **(a)** *Park7*, **(b)** *Nfe2l2*, or **(c)** *Cyp1b1* (n=3). Each red dot represents a DEG (FDR < 0.05) and the total number of up- and downregulated DEGs are indicated. The x-axis represents the log_2_-fold change and y-axis the -log_10_ p-value. **(d-f)** Heatmaps showing the differences in pathway enrichments of downregulated DEGs from **panels a-c** using the **d)** MSigDB Hallmark, **e)** WikiPathway, and **f)** ENCODE project and ChEA databases, focusing on the top 5 most enriched pathways or TFs upon *Park7* depletion. The -log_10_ p-values of the enrichments are depicted as color scale.

Finally, we asked whether our findings in mouse astrocytes are also of relevance for human cellular models of PD. For this we took advantage of recent transcriptomic data of male iPSC-derived astrocytes with *PARK7* mutation (c.192G>C) causing DJ-1 deficiency ^34^ (Mencke *et al.* in preparation). In the human astrocytes, 291 genes were significantly differentially expressed (FDR < 0.05) with 179 of those genes downregulated upon loss of DJ-1 (Fig. 8a). Albeit the number of available genes for enrichment analysis was limited, also in human astrocytes the most enriched pathways were EMT and interferon signaling for down- and upregulated genes, respectively (Fig. 8b-c). Suggesting that many of the male-specific changes upon DJ-1 depletion in mouse could also be relevant for PD diagnosis and disease progression in humans.

**Figure 8.**
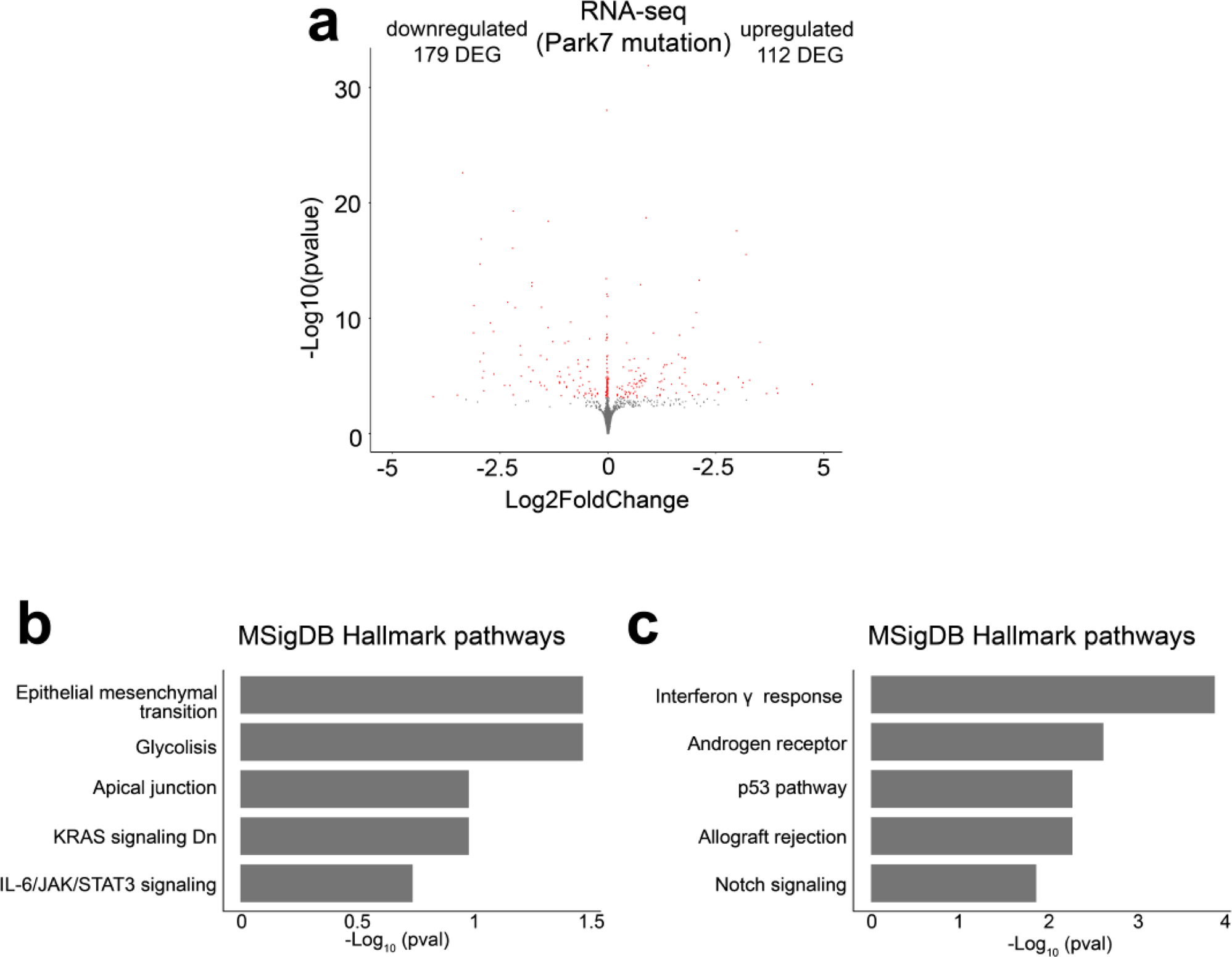
DJ-1-dependent transcriptomic changes in human iPSC-derived astrocytes. **(a)** Volcano plot depicting the DEGs in male iPSC-derived astrocytes with *PARK7* mutation leading to DJ-1-deficiency when compared to isogenic control cells (n=3). Each red dot represents a DEG (FDR < 0.05) and the total number of up- and downregulated DEGs are indicated. The x-axis represents the log_2_-fold change and y-axis the -log_10_ p-value. **(b-c)** Pathway enrichment analysis for the 179 downregulated and 112 upregulated DEGs from **panel a**. The top 5 results from MSigDB Hallmark pathways based on the significance of enrichment are shown for (b) downregulated and (c) upregulated genes.The x-axis represents the -log_10_ p-value of pathway enrichment.

## Discussion

Here, we have characterized the early transcriptome changes occuring in the midbrain of *Park7*^−/−^ mice during aging. For the first time, we report alterations specifically in the male midbrain, in keeping with the epidemiological data on higher PD prevalence in males ^35^. Moreover, many of the transcriptional changes observed in males are affecting genes with sex-specific expression. Consistently, a homozygous female carrier of the c.192G>C mutation of *PARK7* found in our human iPSC line was reported to be clinically asymptomatic at an age when a male patient was already affected ^36^. The gene expression changes in the midbrain of male *Park7^−/−^* mice at 8 months of age were consistent with a downregulation of antioxidant response and reduction in the expression and activity of NRF2. Interestingly, expression of *NFE2L2* and NRF2 target genes have been reported to be decreased in the midbrain of PD patients^37^ and DJ-1 is known to be involved in the stabilization and activation of NRF2^17^. In *Park7*^−/−^ mice, the disruption of DJ-1 function alone might be sufficient to increase reactive oxygen species (ROS) production, but with the parallel decrease in the activity of NRF2 and the transcription of antioxidant genes, the accumulation of ROS is likely to lead to oxidative stress, one of the main hallmarks of PD ^7–9^. Interestingly, these changes seem to take place selectively in the midbrain as no alterations were detected in the cortical tissues of the same male mice, in keeping with midbrain being the most affected area in PD ^6,38^.

It is unclear how the DJ-1-NRF2 interaction occurs. However, it is known that DJ-1 is able to activate PI3K-Akt and ERK pathways to exerts its cytoprotective role in an oxidative stress-associated neurotoxic context model of PD ^39^. Moreover, PI3K-Akt pathway can activate NRF2 signaling ^40,41^ and was found enriched among DEGs in *Park7^−/−^*mice and upon *Park7* knockdown in primary astrocytes. Therefore, in the absence of DJ-1, PI3K-Akt signaling may be reduced (possibly through the inhibition of PTEN ^42,43^), leading to a decreased stabilization of NRF2 and reduced expression of its target genes. Interestingly, a significant decrease in the activation of PI3K-Akt pathway has been observed in the SNpc of PD patients ^43^ and a decline in NRF2 expression has been detected in AD and PD patients with age-dependent cognitive loss ^44–46^.

Our data point to an important role of astrocytes in the gene expression changes and decreased NRF2 signaling induced by loss of *Park7*. Indeed, reduction of NRF2 in astrocytes has been associated with increased PD pathology ^47^, while NRF2 overexpression induces cytoprotection and reduction of symptoms ^44,46,48,49^. Consistently, *Park7* and *Nfe2l2* knockdowns in male astrocytes, but not in microglia or oligodendrocytes, led to downregulation of *Cyp1b1* and *Cdh1* expression, in keeping with the results obtained *in vivo*. Moreover, *Cyp1b1* knockdown alone also decreased expression of *Cdh1* and other target genes, linking CYP1B1 enzyme to the enriched processes such as EMT, focal adhesion, and ECM-interaction in astrocytes. Interestingly, an association of CYP1B1 with the regulation of EMT and redox homeostasis in astrocytes has already been reported ^50^. While astrocytes are unlikely to undergo classical EMT upon loss of *Park7*, astrogliosis has been proposed to be an EMT-like process involving a subset of EMT-related genes, in particular those linked to ECM and adhesion^51^. Remarkably, ECM and focal adhesion were found among the common PD-associated transcriptome signatures in a recent analysis of midbrain neurons derived from PD patients with many different genetic causes ^52^. And finally, DJ-1 has already been associated with alterations in the ECM and promotion of EMT in the context of cancer ^53^.

Importantly, the top upstream regulator predicted to explain expression changes in the midbrain of *Park7^−/−^* male mice at 8 months old was *17β*-oestradiol. Therefore, the neuroprotective role that *17β*-oestradiol, the main female hormone, plays in the brain, including reduction of ROS levels or activation of antioxidant enzymes ^54,55^, may lead to the sexual dimorphism observed in the midbrain transcriptomic profiles of *Park7*^−/−^ mice at 8 months of age ^12^. This is consistent with the fact that incidence and prevalence of PD are higher in males than in females, and that the severity of the symptoms in males are greater ^5,56^. In females, the onset of PD correlates positively with onset of menopause and fertile life span ^57^. Shorter fertile period or lower exposure to estrogen during life are associated with PD in females ^58,59^, while increased endogenous estrogen exposure is associated with reduced severity of motor impairment and later onset of PD ^60^. Interestingly, the menstrual cycle, a period in which the levels of estrogen strongly fluctuate, can influence in the severity of symptoms in females with PD ^61^. Indeed, low-dose estrogen treatment during 8 weeks in Parkinsonian postmenopausal females with motor fluctuations had a significant decrease in motor disability ^62^.

Thus, there is an increasing body of evidence for sex-specific differences in the phenotypic presentation of Parkinson’s disease (PD) that also include specific changes in gene expression ^63^. As PD is a recognized as a heterogenous disorder, monogenic forms of PD represent prototypes for dissecting subtypes of PD with different pathophysiological mechanisms involved. DJ-1 is a rare form of early-onset PD with only few families affected world-wide, so that only limited clinical observations are available. Among these the authors characterized a family with a striking difference in disease expression between the two homozygous mutation carriers ^64^: whereas the male carrier presented with a typical early onset PD starting at age 35, the sister was still presenting no motor symptoms at age 42 and still normal PET Scan results.

Also *all-trans* retinoic acid appeared among the top upstream regulators linked to the gene expression changes in male *Park7*^−/−^ mice. Interestingly, *17β*-oestradiol and *all-trans* retinoic acid, after binding to and activating their receptors (estrogen receptors (ERs) and retinoic acid receptors (RARs), respectively), can alter the transcription of their target genes, many of which are shared, exerting opposing effects through genomic antagonism ^28^. Some of the known shared target genes between EREs and RARs are related with EMT, ECM, focal adhesion, and antioxidant response ^28,65–68^. Therefore it is interesting that CYP1B1 is involved in the metabolism of both *17β*-oestradiol and *all-trans* retinoic acid^69^. Importantly, *Cyp1b1* is among the NRF2-target genes ^32^, suggesting that NRF2, via transcriptional modulation of *Cyp1b1*, may alter *17β*-oestradiol and *all-trans* retinoic acid levels, with consequences on neuroprotection and regulation of ECM, EMT and focal adhesion, as discussed above.

Finally, another potential upstream regulator we identified was TGF-β. The activity of TGF-β superfamily can be activated by ROS ^70^. Therefore, upon an increase in ROS levels in the midbrain, TGF-β signaling could be extraordinarily activated, promoting EMT-related processes. There is a number of reports indicating an interaction between NRF2 signaling and TGF-β pathway via SMAD effector proteins ^71–73^ Importantly, TGF-β signaling is one of the main pathways regulating GFAP promoter activity and expression ^74^ and astrocyte-derived TGF-β is involved in the crosstalk between astrocytes and the rest of brain cells, modulating their functions ^70^. Therefore, TGF-β signaling is an important regulator of astrocyte formation and function, and its disruption in astrocytes has been associated with pathogenesis of neurological disorders, including PD ^75,76^.

Importantly, transcriptomic changes in human male iPSC-derived astrocytes with *PARK7* mutation showed overlap regarding the affected pathways with those found *in vivo* in mice, including EMT as the most enriched pathway for the downregulated genes. This supports the relevance of our findings for understanding PD progression and diagnosis in humans. In addition, upregulated genes were enriched for interferon signaling, similarly to the *Park7*-depleted mouse astrocytes. Interestingly, interferon signaling regulates the activation of PI3-Akt pathway that, as discussed above, may also be altered in the astrocytes of *Park7*^−/−^ mouse midbrain ^65^.

Taken together, we have described an age- and sex-specific transcriptome signature induced by loss of *Park7* and reveal it to involve dysregulation of EMT, ECM, and focal adhesion associated genes by NRF2 and CYP1B1 in midbrain astrocytes. These results open new avenues for investigation of mechanisms underlying PD progression and its sexual dimorphism.

## Material and Methods

### Animals

Mouse colonies were bred and maintained by the Animal Facility of the University of Luxembourg, on a 12 hour light/dark cycle with provided food and water *ad libitum*. All the experiments were conducted following the national guidelines of the animal welfare law in Luxembourg (Règlement grand-ducal adopted on January 11th, 2013), which was approved by the Animal Experimentation Ethics Committee (AEEC) and that follows the European Union guidelines (2010/63/EU).

### Park7^−/−^ mice

The *Park7^−/−^* mice (B6.129P2-Park7Gt (XE726) Byg/Mmucd) used in the present study were already previously characterized ^22^. The mice had been backcrossed with C57BL/6N strain for at least ten generations. The *Park7*^−/−^ and *Park7*^+/+^ mice used in our studies were sex- and age-matched siblings generated from heterozygous *Park7*^+/−^ breeding pairs. The *Park7*^−/−^ and *Park7*^+/+^ mice were bred in-house during three and eight months, to generate two different study cohorts. For each cohort, both males and females were used. A total of 8 mice (4 male and 4 female) per genotype was used at 3 months of age, while 12 mice (8 male and 4 female) per genotype were used at 8 months of age. Mice were anesthetized with a mixture of ketamine and medetomidine (150 and 1 mg/kg, respectively) and transcardially perfused with PBS (phosphate-buffered saline). After that, brains were extracted and split along the longitudinal fissure in two hemi brains, as described in Karunakaran et al. 2007 ^77^. Ventral midbrains and cortices of both hemispheres were dissected, snap-frozen in dry ice and stored at −80°C until use.

### iPSC-derived astrocytes

The used isogenic iPSC lines C4 and C4mut, harboring the pathogenic c.192G<C mutation, were previously published ^78^. Astrocytes were generated from smNPC via hNSCs as described in Palm et al. 2015^79^. 2 days prior to hNSC differentiation, 400 k smNPCs were seeded into one 6-well of a 6-well plate per line. After 2 days, the medium was changed to smNPC medium with 20 ngng/ml FGF-2 (Peprotech - 100-18B). After 4 days, the cells were split with Accutase® (Sigma A6964) and the medium was changed to hNSC medium consisting of DMEM/F12 w/o HEPES (Life/Tech – 21331046) supplemented with N2 supplement (Life/Tech – 17502048), B27 supplement with vitamin A (Life Technologies Europe BV/Thermo Fisher Scientific 17504044), GlutaMAX Supplement (Life/Tech – 35050-061), penicillin/streptomycin (Life/Tech – 15140-163), 40 ng/ml EGF (Peprotech - AF-100-15-1mg), 40 ng/ml FGF-2 (Peprotech - 100-18B) and 1.5 ng/ml hLIF (Peprotech - AF-300-05). hNSCs were split with Accutase® when reaching 70-80 % of confluence. The astrocytic differentiation medium consisted of the basic cultivation medium DMEM/F12 w/o HEPES (Life/Tech – 21331046) supplemented with 1 % penicillin/streptomycin (Life/Tech – 15140-163), 1 % GlutaMAX Supplement (Life/Tech – 35050-061) and 1 % fetal bovine serum (Life/Tech – 10270-106). 1 million hNSCs per T25 flask were plated 2 days prior to astrocyte differentiation. After 2 days, hNSC medium was changed to astrocyte medium. After 40 days, astrocytes were split to get rid of neurons that are dying during the differentiation and during the process of splitting. After 60 days, astrocytes were considered to be mature and all experiments were conducted around day 60.

### Primary cultures and cell isolation

Mixed glial cell cultures were obtained from the brains of 1-3 days old newborn CD1 mice (Charles River, France). After removing meninges and large blood vessels, brains were mechanically dissociated in PBS (phosphate-buffered saline). To keep males and females separated, each individual brain was plated in one T75 flask with culture medium composed of DMEM (Dulbecco’s Modified Eagle Medium), 10% FBS (fetal bovine serum) (Life Technologies, 10270-106), 100 U/ml penicillin and 100 μg/ml streptomycin. Furthermore, a small piece of tail (2-3 mm) from each newborn mouse was retained in order to subsequently perform individual sex genotyping (see next section). Mixed glial cell cultures grew at 37°C in a 5% CO2 humidified atmosphere and the medium was changed twice a week.

Two weeks later, once cultures were confluent, cell isolation was performed using a magnetic cell sorting (MACS) method following the manufacturer’s instructions (Miltenyi Biotec, The Netherlands). Isolation of specific cell types was performed separately in male and female cultures An anti-CD11b microbeads antibody (Miltenyi Biotec, 130-049-601) and an anti-O4 microbeads antibody were used to isolate microglia and oligodendrocytes, respectively (positive selection for both). For astrocytes, the negative fraction was kept. To have enough cells, microglia of the same sex were mixed and plated in culture medium composed by mixed glial cell culture-conditioned medium and DMEM (50/50, v/v). O4^+^ cells of the same sex were also mixed and kept in culture medium composed by MACS Neuro Medium (Miltenyi Biotec, 130-093-570), 2% MACS NeuroBrew-21 (1x) (Miltenyi Biotec, 130-093-566), 0.5 mM L-Glutamine (Lonza, #BE17-605E), 100 U/ml penicillin and 100 μg/ml streptomycin, 10 ng/mL Platelet Derived Growth Factor AA (PDGF-AA) (Miltenyi Biotec, 130-093-978) and 10 ng/mL Fibroblast Growth Factor 2 (FGF-2) (Miltenyi Biotec, 130-105-787), for 14 days with medium change every 2 days. Astrocytes of the same sex are also mixed and kept in culture medium composed of DMEM, 10% FBS, 100 U/ml penicillin and 100 μg/ml streptomycin for 2 weeks, with a second negative selection in between (after 7 days).

### Genotyping for sex of primary cultures

Tail fragments (2-3 mm) from each newborn mouse were placed in 1.5 ml eppendorf tubes. After adding 200 μl of Direct PCR Tail (Lysis buffer) (Viagen, 102-T) and 4 μl of proteinase K (Invitrogen, 25530-049), samples were kept for at least 5 hours at 55°C with occasional vortexing, to extract the genomic DNA. Afterwards, an equal volume of isopropanol was added, followed by vortexing and centrifugation at 13 000 rpm at 4°C for 10 minutes. DNA was washed with 200 μl of 70% ethanol and resuspended in 350 μl of DNase/RNase-free water. Afterwards, PCR was performed to amplify the genes *Uba1* and *Ube1y1* on the X and Y chromosome, respectively. For this, we used *Ube* primers that amplify 2 products in males and only the larger product in females ^80^. PCR sex genotyping reactions were performed in a final volume of 25 μl with the following reagents and volumes per sample: 12.5 μl KAPA2G Fast HotStart DNA Polymerase (5 U/μL) from the KAPA2G Fast HotStart PCR Kit (Sigma-Aldrich, KK5601), 1 μl of 12.5 μM *Ube* forward primer and 1 μl of 12.5 μM *Ube* reserve primer, 9.5 μl dH2O and 1 μl of purified gDNA; and the following PCR parameters: initial denaturation at 94°C for 2 min, 35 cycles with 94°C for 30 s, 57°C for 30 s, and 72°C for 30 s, followed by final elongation at 72°C for 5 min. PCR products were analyzed using agarose electrophoresis together with a DNA ladder (Invitrogen, 10488-058) on 2% agarose gels that contains SYBR safe (Invitrogen, G601802) for the visualization under UV-illumination.

### RNA interference in primary cultures

Knockdown of the *Park7*, *Nfe2l2,* and *Cyp1b1* in primary cultures was done using gene-targeted siRNAs. Primary astrocytes were seeded at a density of 4×10^5^ cells/well into 12-well plates, while primary microglial cells and oligodendrocytes at a density of 3×10^5^ cells/well into 24-well plates. Later, microglia, astrocytes and oligodendrocytes were transfected 2 days, 7 days and 14 days after isolation, respectively. Transfections of 30 nM of siRNAs targeting *Park7*, *Nfe2l2*, *Cyp1b1* or a negative control siRNA (Eurogentec, Belgium) were done using Lipofectamine RNAiMAX reagent (Life Technologies, Belgium) for 24 hours. siRNAs and Lipofectamine RNAiMAX were prepared in DMEM for astrocytes and microglia, while for oligodendrocytes they were prepared in the culture medium (MACS Neuro Medium, 2% MACS NeuroBrew-21 (1x), 0.5 mM L-Glutamine (Lonza), 100 U/ml penicillin and 100 μg/ml streptomycin, 10 ng/mL PDGF-AA and 10 ng/mL FGF-2). After 24 hours, RNA from the different cells was extracted.

### RNA extraction

For all mouse midbrain samples, RNeasy® Plus Universal Mini Kit (Qiagen, Germany) was used to extract the total RNA. For the different primary cultures, innuPREP RNA Kit (Westburg, The Netherlands) was used to extract the total RNA after 24 hours of siRNA transfection. Total RNA was extracted from iPSC derived astrocytes using the Qiagen RNeasy Kit according to manufacturer’s protocol.

### RT-qPCR

For cDNA synthesis, depending on the sample availability, 100 ng, 300 ng, or 500 ng of total RNA were used. Total RNA was mixed with the following reagents to get a final volume of 40 μl: dNTPs (0.5 mM, ThermoFisher, R0181), oligo dT-primer (2.5 μM), 1 μl RevertAid reverse transcriptase (200 U/μl, ThermoFisher, EP0441), and 1 μl Ribolock RNase inhibitor (40 U/μl, ThermoFisher, EO0381). The reverse transcription was performed at 42°C for 1 hour and stopped by incubating at 70°C for 10 min. cDNA was diluted 1:5 or 1:3 in DNase/RNase-free water, in the case of using 500 ng or 300 ng of total RNA, respectively.

qPCR reactions were performed using 5 μl of diluted cDNA, 5 μl of a mix containing 2 μM custom designed primers and 10 μl of 1xAbsolute Blue qPCR SYBR green low ROX mix (ThermoFisher, AB4322B), in a final volume of 20 μl per well. Specificity of the primers was verified previously by detecting a single melting point in the melt curve. qPCR parameters were the following: 95°C for 15 min and then 40 cycles of 95°C for 15 sec, 55°C for 30 sec and 72°C for 30 sec. Gene expression levels were normalized following the 2^-(ΔΔCt)^ method. Briefly, the following formula (ΔCt(_target gene_)-ΔCt(_housekeeping gene_))_test condition_ - (ΔCt(_target gene_)-ΔCt (_housekeeping gene_))_reference condition_ was used to get ΔΔCt. *Rpl13a* was used as the housekeeping gene. To perform statistics, we used GraphPad Prism 9 and to determine the significance, unpaired t-test was applied to *Park7^−/−^* midbrain and cortex samples, and one sample t-test was applied in the primary cultures.

### RNA-seq

RNA was extracted as described above and its quality was determined by using the Agilent RNA 6000 Nano kit (5064-1511) in an Agilent 2100 Bioanalyzer machine. All samples had a RIN value above 7. mRNA sequencing was performed by the Sequencing Platform of the Luxembourg Centre for Systems Biomedicine (LCSB) of the University of Luxembourg. The TruSeq Stranded mRNA Library Prep kit (Illumina) was used for library preparation. Single-end, stranded sequencing was executed in midbrain samples using an Illumina NextSeq 500 machine with a read length of 75 bp, while paired-end, stranded sequencing was executed in primary astrocytes using a NextSeq2000 machine with a read length of 50 bp.

### RNA-seq data analysis

FastQC (v0.11.9) was used to assess the quality control of the raw reads ^81^. Reads were trimmed by AdapterRemoval (v2.3.1) ^82^. SortMeRNA (v2.1) was used to remove rRNA reads ^83^, followed by mapping using STAR (v.2.7.4a) ^84^. The mouse reference genome used was GRCm38 release 102. Picard tool (v2.10.9) ^85^ was used to validate the BAM files. After getting the BAM files, featureCounts from the R package Rsubread (v1.28.1) ^86^ produced the counts of the reads. Using the counts as input, we used R package DESeq2 (v1.20.0) ^87^ to perform the differential expression analysis and obtain the differentially expressed genes (DEGs). RUVSeq (v1.20.0) was used to remove unwanted variation (batch correction). For primary astrocytes the DEGs were identified using paired analysis due to high variation between between cell isolations. Enrichment analysis for DEGs was conducted with EnrichR ^88–90^ and Ingenuity Pathway Analysis (IPA) ^91^. The distribution of the per-gene dispersion was estimated using DESeq2. The dispersion estimates were obtained after sub-setting the main DESeq2 object with all samples using the different conditions of interest, and then merged into a single table for plotting boxplots and violin plots.

### Immunocytochemistry

Mix of glial cells after 14 days in culture, astrocytes after 2 weeks in culture, microglia after 48 hours in culture and O4 positive cells (pre-oligodendrocytes and mature oligodendrocytes) after 6 days and 14 days in culture, were seeded on poly-L-lysine coated coverslips into 24-well plates at a density of 3×10^5^ cells/well ^92^ and fixed with 4% PFA for 20 min at room temperature. For permeabilization, 0.3% Triton X-100 in PBS was used for 5 min, followed by 3 washing steps with PBS. Blocking was performed with 3% (w/v) BSA for 30 min at room temperature, followed by an incubation with primary antibodies overnight at 4°C. Mix of glial cells were co-stained using a mouse anti-GFAP antibody (1/300, Cell Signaling, 3670) and a rat anti-F4/80 antibody (1/300, Bio-Rad, MCA497) to check astrocytic and microglial populations. Astrocytes and microglia cultures were also co-stained with the same anti-GFAP and anti-F4/80 antibodies, to check the purity of those cultures and discard any other cell type contamination. Mouse anti-O4 (1/100, R&DSystems, MAB1326) and rat anti-MBP (1/50, Abcam, ab7349) antibodies were also used to check for pre-oligodendrocytes and mature oligodendrocytes respectively. The next morning, cells were incubated with secondary antibodies: Alexa Fluor 488-conjugated donkey anti-mouse (1/1000, ThermoFisher Scientific), Alexa Fluor 488-conjugated donkey anti-rat (1/1000, ThermoFisher Scientific), Alexa Fluor 568-conjugated goat anti-rat (1/1000, ThermoFisher Scientific), or an Alexa Fluor 568-conjugated donkey anti-mouse (1/1000, ThermoFisher Scientific) for one hour at room temperature. Glass slides with the cells were washed with PBS and mounted with DAPI Fluoromount-G (SouthernBiotech, USA). LSM 510 META inverted confocal microscope at a 40× magnification (Carl Zeiss Micro Imaging, Gottingen, Germany) was used to take the pictures.

### Primers

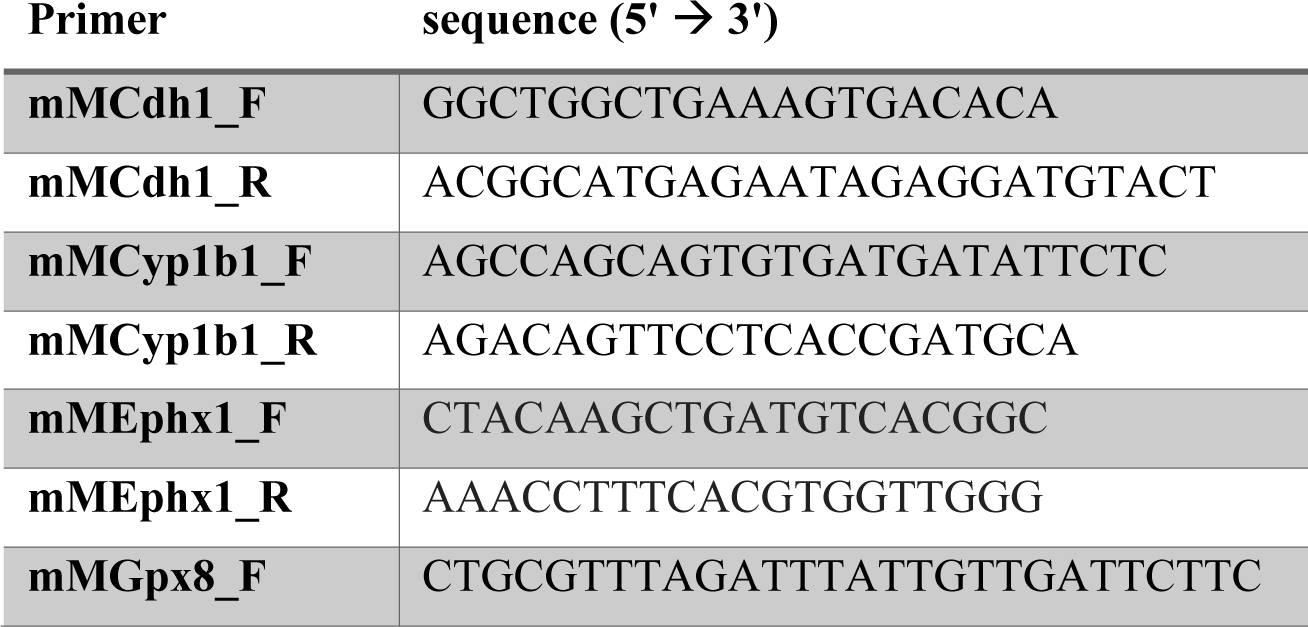
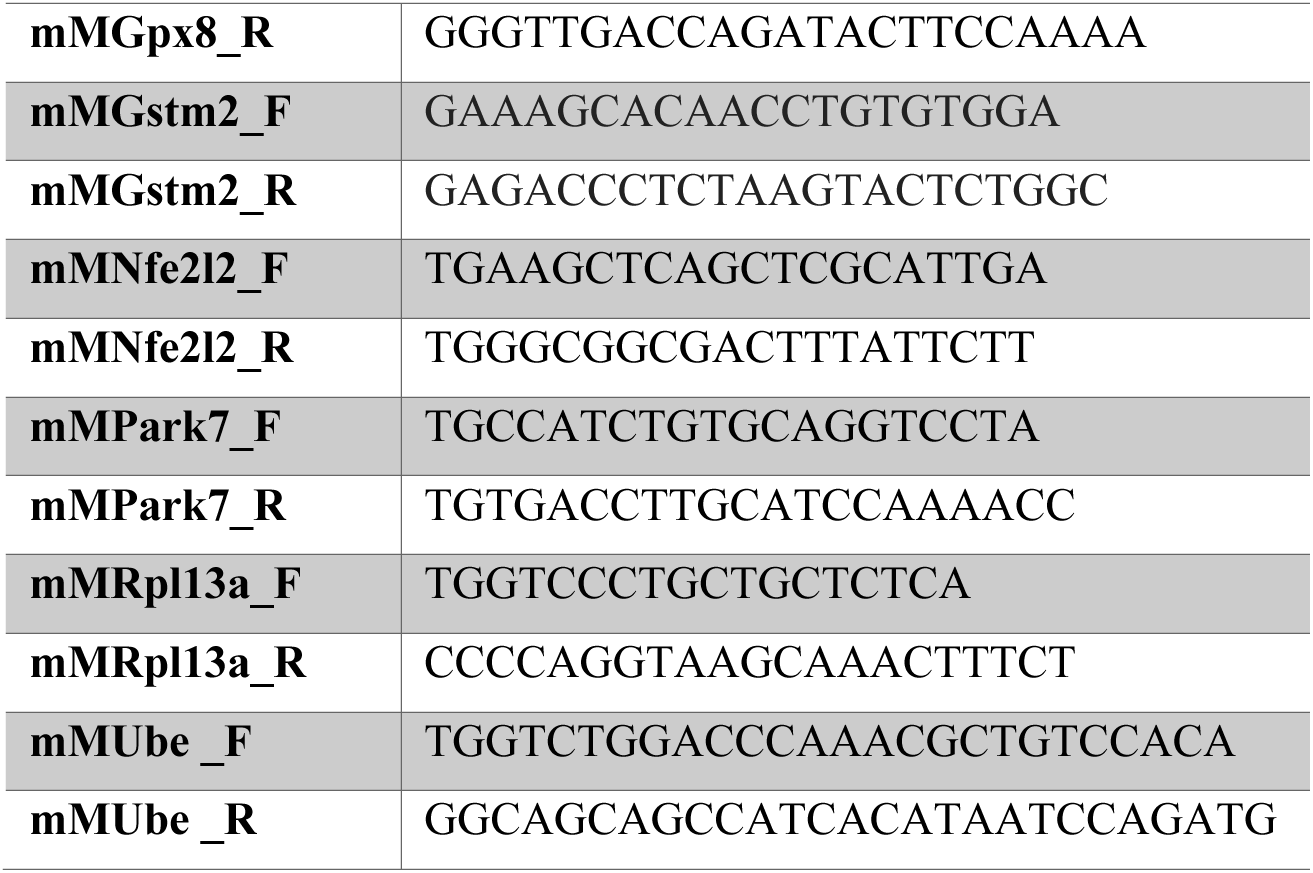

### siRNAs

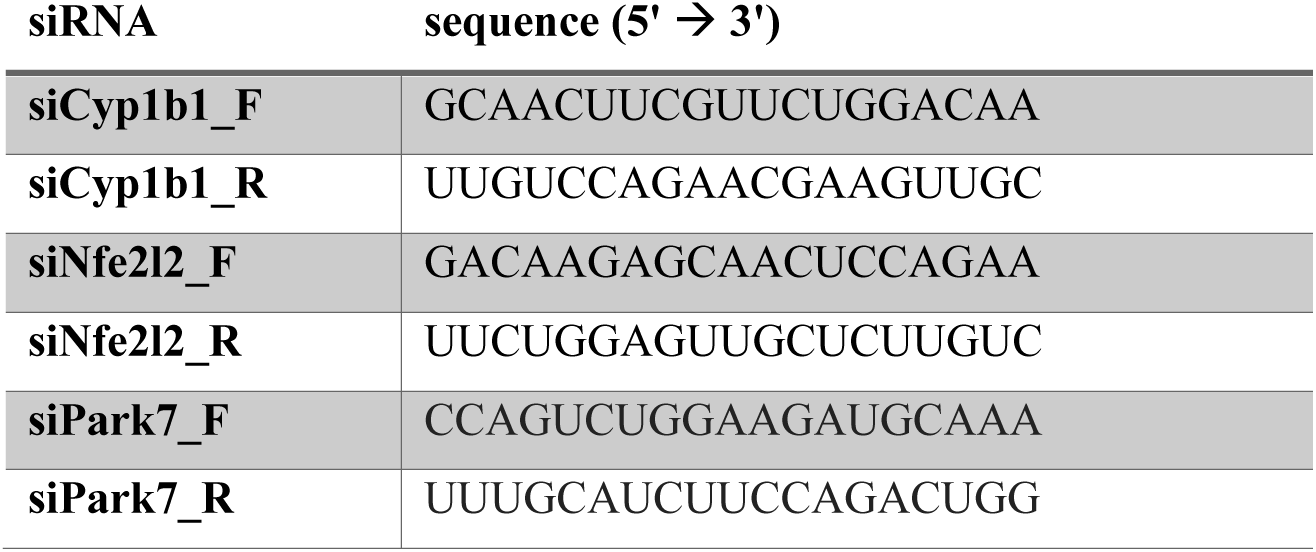

## Author Contributions

MB and LS conceived the project with input from MM and TS. SH, TH, AS, MB, and LS designed the experiments and analysis. SH, AS, PG and MB prepared mouse tissues. SH performed all mouse RNA-seq experiments and most bioinformatic analysis. SH and TH prepared mouse cell cultures and performed RNAi experiments. TH performed immunostainings. PM, IB and JO performed the RNA-seq experiments and bioinformatic analysis of the human iPSC-derived astrocytes. RH prepared the libraries and performed the sequencing. YG supported bioinformatic analysis and developed methodology. SH, TH, AS, MB, MM, TS and LS analyzed and interpreted data. RK, MM, MB, and LS supervised the project. SH and LS wrote the manuscript. All authors read and approved the final manuscript.

## Acknowledgments

We would like to thank Drs Aurélien Ginolhac and Anthoula Gaigneaux for their support with bioinformatic analysis and and Dr Djalil Coowar (Animal Facility of University of Luxembourg) for help with breeding of experimental mice. The computational analysis presented in this paper were carried out using the HPC facilities of the University of Luxembourg.

## Data and code availability

The RNA-seq data for this study have been deposited in the European Nucleotide Archive (ENA) at EMBL-EBI under accession number PRJEB59115 (https://www.ebi.ac.uk/ena/browser/view/ PRJEB59115).

All the RNA-seq analysis code can be found in the following repository: https://github.com/sysbiolux/Helgueta_et_al_2023

