## Supplementary Figures and Legends, Supplementary Table Legends for "*Park7* deletion leads to age- and sex-specific transcriptome changes involving NRF2-CYP1B1 axis in mouse midbrain astrocytes"

**Supplementary Information**

**Supplementary Figures**

**Supplementary Table Legends**

**Supplementary Information**

The Supplementary Information contains four Supplementary Figures and five Supplementary Tables.

**Supplementary Figure 1. Distribution of the per-gene dispersion in *Park7$^{-/-}$* mouse midbrain.** Per-gene dispersions to estimate the median dispersion each sample set show similar results in all sample sets, except for the second cohort of 8-month-old male samples. Lack of changes in 3 and 8-month-old females are demonstrated to not be due to technical or biological (oestrus cycle) variation between the female samples. Increased dispersion in the second cohort of 8-month-old male samples partially explains the lower number of DEGs compared to the first cohort.

**Supplementary Figure 2. Loss of *Park7* in female mice does not lead to gene expression changes associated with NRF2 signaling, epithelial-to-mesenchymal transition, focal adhesion, and extracellular matrix composition.** (a-d) Pathway enrichment analysis for the 101 DEGs altered in 8-month-old female mice. The top 5 results from (a) KEGG pathways, (b) WikiPathway pathways, (c) MSigDB Hallmark pathways, and (d) GO terms for cellular components based on the significance of enrichment are shown.The x-axis represents the -log10 p-value of pathway enrichment. Complete results of enrichment analysis are available in the Supplementary Table S4. (e) Top 5 TFs associated with DEGs from Park7-/- female mice based on the significance of enrichment for primary TF targets from the ENCODE project and ChEA database. The x-axis represents the -log10 p-value. (f) Top5 Upstream Regulators associated with DEGs from Park7-/- female mice predicted by Ingenuity Pathway Analysis (IPA). The x-axis represents the -log10 p-value of pathway enrichment.

**Supplementary Figure 3 Genotyping of newborn mice prior to glial cell isolation.** *Ube* primers were used to test whether DNA from newborn mouse tails carried XY (two separate bands) or XX (one stronger band) genotype. L = Ladder.

**Supplementary Figure 4. Enrichment analysis of upregulated genes upon *Park7* knock-down. (a-c)** Heatmaps showing the differences in pathway enrichments of upregulated DEGs from **panels a-c** using the **(a)** ENCODE project and ChEA databases, **(b)** BioPlanet, and **(c)** MSigDB Hallmark databases, focusing on the top 5 most enriched pathways or TFs upon *Park7* depletion. The -$\log_{10}$ p-values of the enrichments are depicted as color scale.

**Supplementary Table 1. List of DEGs from *Park7*$^{-/-}$ mice compared to wild type littermate controls.** The separate worksheets include DEGs from 3-month-old female mice, 3-month-old male mice, 8-month-old female mice, 8-month-old male mice (first cohort), 8-month-old male mice (second cohort), 8-month-old male mice (first and second cohort combined using batch correction), with FDR < 0.05. The table includes ENSEMBL IDs, base mean, $log_2$-fold change, p-value, FDR (padj), and gene symbol for each DEG.

**Supplementary Table 2. Mapped reads for the 4 groups of mice and the 3 groups of astrocytes.**

**Supplementary Table 3. Lists of enriched pathways in males.** Separate worksheets include enrichments for analysis using DEGs obtained from the comparison of 8-month-old male *Park7*$^{-/-}$ mice with the wild type littermate controls (first and second cohort together) from KEGG, WikiPathway, MSigDB, and BioPlanet databases, and the GO terms for Cellular components.

**Supplementary Table 4. Lists of enriched pathways in females.** Separate worksheets include enrichments for analysis using DEGs obtained from the comparison of 8-month-old female *Park7*$^{-/-}$ mice with the wild type littermate controls from KEGG, WikiPathway, MSigDB, and BioPlanet databases, and the GO terms for Cellular components.

**Supplementary Table 5. Lists DEGs in primary mouse astrocytes upon the knock-down of *Park7*, *Nfe2l2*, or *Cyp1b1*.** DEGs were defined by applying FDR < 0.05. The table includes ENSEMBL IDs, base mean, $log_2$-fold change, p-value, FDR (padj), and gene symbol for each DEG.

### Supplementary Figure 1

**a**

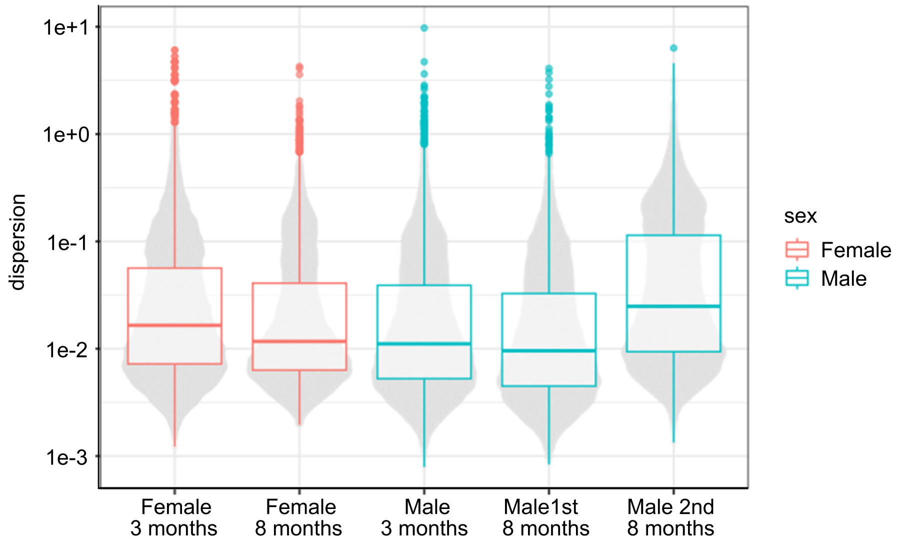

### Supplementary Figure 2

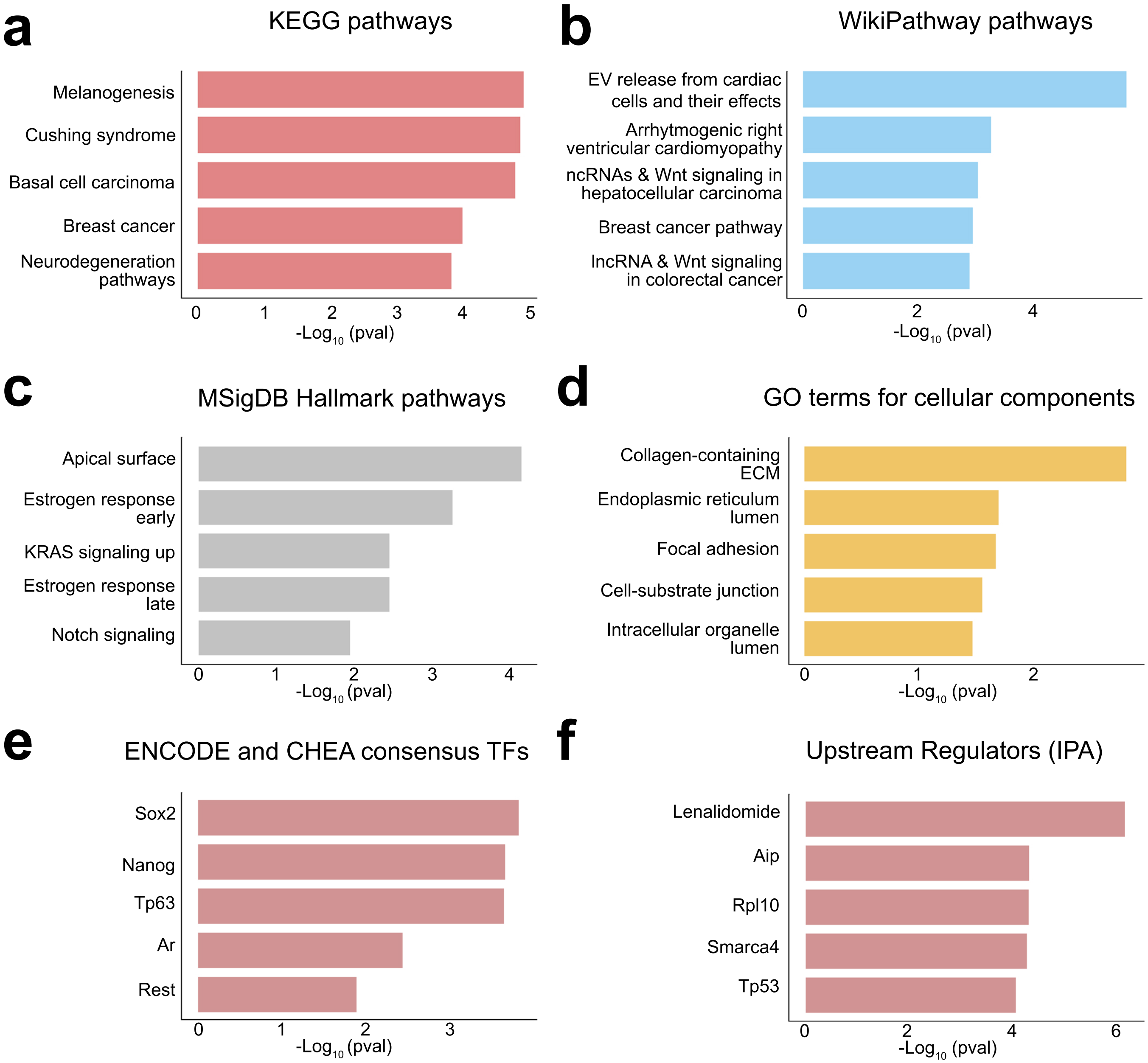

### Supplementary Figure 3

a

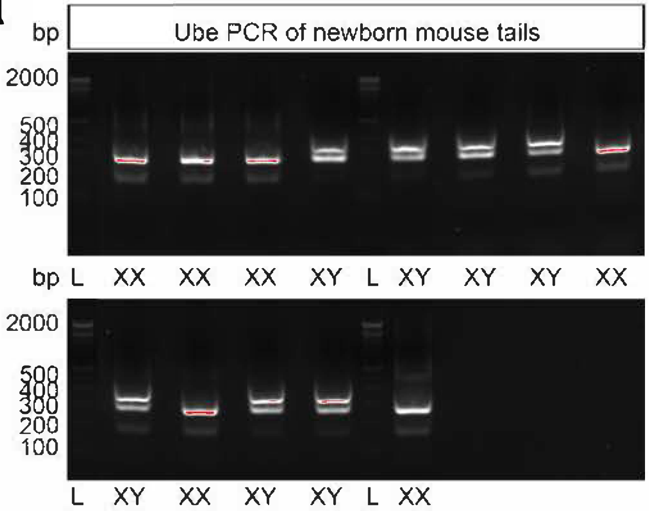

### Supplementary Figure 4

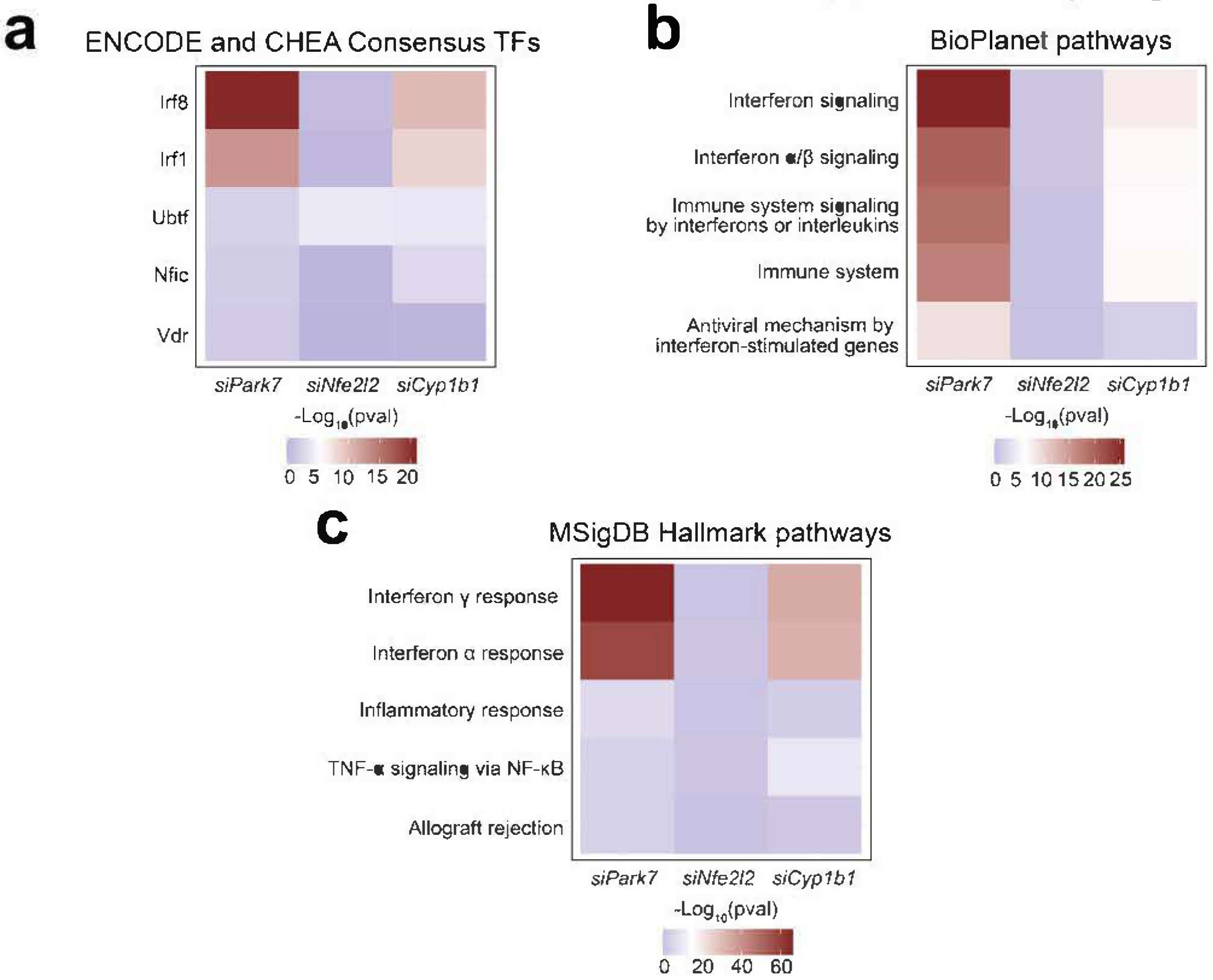
